# A medial entorhinal connectome reveals compartmentalized microcircuit architecture

**DOI:** 10.64898/2026.10.28.762527

**Authors:** Te Shi, Yinda Chen, Haiqun Jin, Haitao Jiang, Chunying Yin, Jun Guo, Haoyuan Shi, Xin Ren, Yunjie Liao, Yirui Zhang, Ankang Lu, Jinpeng Lu, Hao Xie, Tonghuan Xiao, Jiateng Shou, Peiyao Shi, Wenlin Jin, Haoxing Zhu, Cheng Li, Xuejin Chen, Zhiwei Xiong, Feng Wu, Ruobing Zhang

## Abstract

The medial entorhinal cortex (MEC) transforms multimodal inputs into spatial representations, but the synaptic architecture underlying its computations remains unclear. Here we generated a synapse-resolution connectomic dataset spanning all six layers of mouse MEC and the adjacent parasubiculum with serial electron microscopy, providing a continuous volume for mapping cellular organization and connectivity across layers and regions. We found that clustered pyramidal neurons in MEC layer II extended oblique apical dendrites with a dorsolateral bias and long-range myelinated axons within the layer. Exhaustive reconstruction of one cluster revealed a central exclusion core (ExC) that spatially partitioned basal dendritic architecture and synaptic connectivity. A spatially constrained growth model reproduced the position-dependent dendritic patterns. Within the ExC, pyramidal neurons formed a compact recurrent excitatory network and converged onto shared inhibitory partners. Basket cells (BC) and recurrent Martinotti cells provided perisomatic and apical-dendritic feedback, respectively. Dense, compact reciprocal connectivity between L2P neurons and BCs within the ExC, together with BC axonal bands extending beyond the ExC, suggests a potential circuit mechanism for generating temporally structured inhibitory output from the pyramidal-cell cluster along directionally extended axons. Together, these findings link cellular-scale geometry and synaptic wiring to emergent circuit dynamics that extend from dense computation within individual clusters to region-wide modulation, providing a connectomic foundation for understanding the circuit basis of spatial navigation.

## Main

The medial entorhinal cortex (MEC) is central to mammalian spatial navigation and contains multiple classes of neurons involved in spatial representation and self-motion processing [1, 2]. Among them, grid cells provide a metric for spatial coding through periodic firing fields that tile the environment [3]. Models of spatial computation emphasize structured recurrent interactions [4, 5], yet how these interactions are embedded in the cellular architecture of the MEC remains unclear.

Electrophysiological recordings and calcium imaging have revealed diverse spatially tuned populations in the MEC, including grid, head-direction and border cells [1, 2]. These functional populations are embedded in a layered and spatially patterned cellular architecture. In layer II, spatially tuned cells occur in both stellate and pyramidal populations [6–10]. Electrophysiological studies have identified selective excitatory connections [11, 12] and recurrent inhibitory circuits [13, 14] in superficial MEC. Anatomical studies further show that layer II pyramidal (L2P) neurons form regularly arranged patches across mammalian species [15–17]. These patches are preferentially targeted by the parasubiculum (PaS) [18], while two-photon imaging reveals grid-cell clusters lying largely in the intervening spaces [19]. How synaptic interactions within and between spatially organized neuronal populations support the precise spatiotemporal activity patterns underlying navigation and memory remains unresolved.

Advances in volume electron microscopy (EM) and computational reconstruction have enabled dense mapping of neuronal morphology and synaptic connectivity within continuous tissue volumes [20, 21]. Large-scale connectomic studies have uncovered cell-type-specific wiring patterns and circuit motifs, revealing organizational principles that link synaptic connectivity to neural computation [22–30]. In rat MEC, dense EM reconstruction revealed path-length-dependent synapse sorting along excitatory axons and a feedforward inhibitory circuit involving myelinated interneuron axons [31]. Understanding how local wiring motifs contribute to the broader organization of MEC circuits requires continuous anatomical maps spanning diverse cell types, cortical layers and neighbouring regions. Such maps can reveal how neuronal compartments are organized and interconnected, establishing a structural basis for investigating the computations that support spatial navigation and memory.

Here we generated a synapse-resolution connectomic dataset spanning approximately 0.56 mm^3^ of mouse brain tissue, encompassing all six MEC layers and the adjacent PaS. Neuronal reconstruction and classification delineated cellular and laminar organization across the volume. Exhaustive reconstruction of one fully contained L2P cluster revealed a central exclusion core (ExC) that partitions the cluster into spatial compartments with distinct basal dendritic architectures and connectivity patterns. Within the core, pyramidal neurons formed a compact recurrent excitatory network with distinct perisomatic and apical-dendritic inhibitory feedback pathways. Models of dendritic growth and spiking activity linked this architecture to position-dependent dendritic patterns and the temporal coordination of recurrent activity. Together, these findings reveal spatial organizing principles of entorhinal microcircuits and provide a structural foundation for investigating the network mechanisms underlying spatial navigation and memory.

## A serial-section electron microscopy dataset of mouse MEC

A tissue volume (Fig. 1a) containing the mouse MEC and adjacent PaS was imaged using automated tape-collecting ultramicrotomy combined with multibeam scanning electron microscopy (ATUM–SEM; Zeiss MultiSEM 505), following established large-volume EM workflows [20] (Fig. 1b–d). The dataset was acquired from 17,913 serial sections, each with a section thickness of 35 nm. After excluding damaged and incomplete sections, 16,833 sections were retained for reconstruction. Images were acquired at 4 nm per pixel in the *x* and *y* dimensions, yielding an imaging voxel size of 4 nm *×* 4 nm *×* 35 nm. The raw acquisition (Fig. 1a, b) covered a volume of 1.036 mm *×* 0.923 mm *×* 0.589 mm (*∼*0.563 mm^3^), spanning all six layers of the medial entorhinal cortex (MEC) and the adjacent parasubiculum (PaS) (Fig. 1e). The complete raw image dataset comprised approximately 1.2 PB.

**Figure 1.**
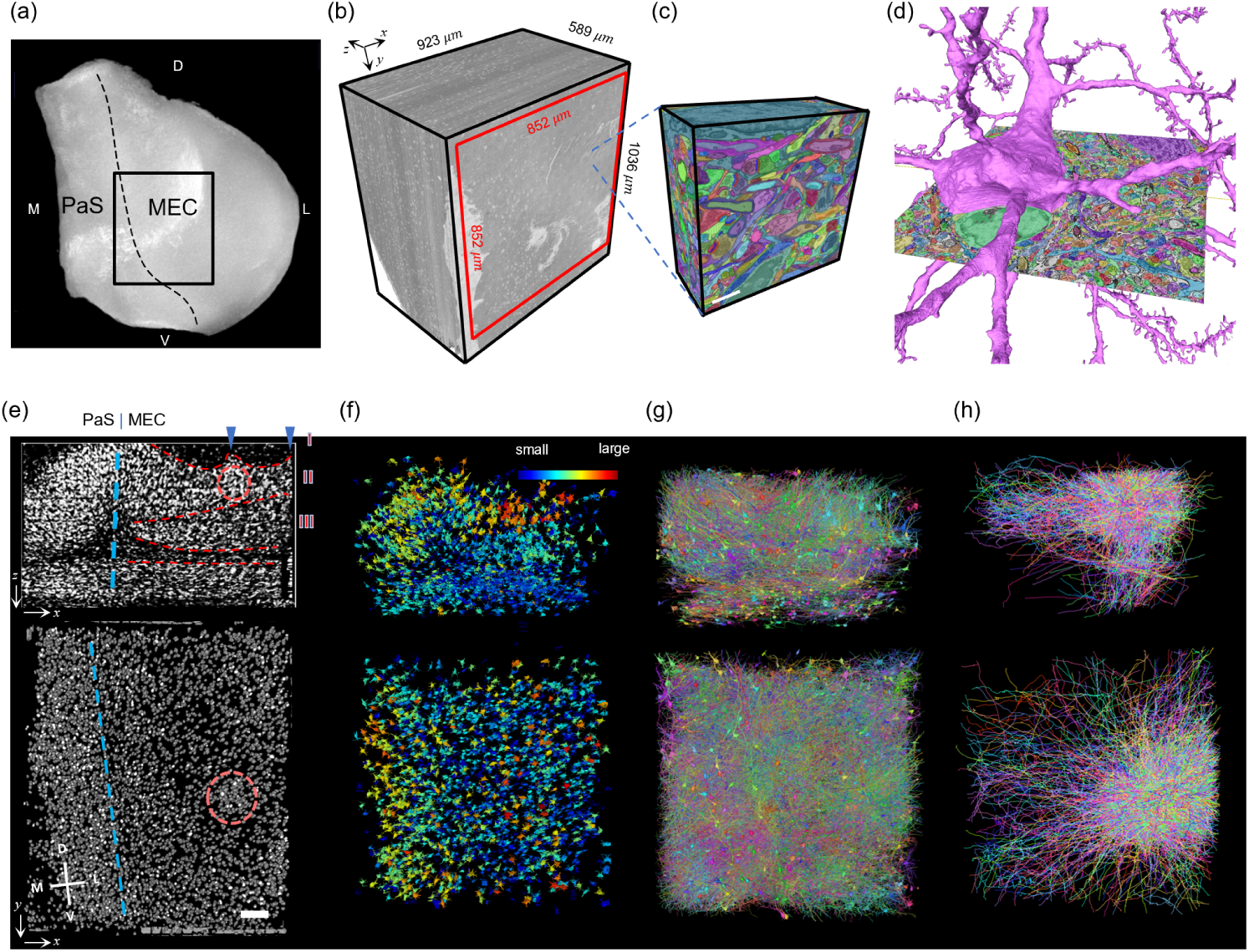
Large-scale EM reconstruction of the mouse medial entorhinal cortex (MEC) and adjacent parasubiculum (PaS). (a) The MicroCT rendering of the MEC sample with the target imaging region indicated by the black box (1.036 mm *×* 0.923 mm). (b) Three-dimensional rendering of the MEC EM volume, spanning 1.036 mm *×* 0.923 mm *×* 0.589 mm (*∼*0.563 mm^3^). (c) Cell and synapse segmentation in a small subvolume, with paired pre- and postsynaptic sites colored red and blue, respectively. (d) A rendering of a pyramidal cell mesh overlaid on the segmentation image. (e) Nuclear distribution in orthogonal horizontal plane (top, *xz*) and tangential plane (bottom, *xy*) sections showing the PaS and layers of the MEC, with a layer II pyramidal cell cluster indicated by a red circle. (f) Example distribution of neuronal somata, with nuclei size represented by color, from a randomly selected subset of neurons. (g) Three-dimensional mesh visualization of randomly selected reconstructed neurons, each assigned with random colors. (h) Demonstration of manually proofread dendritic and axonal skeletons from 150 example neurons around a layer II pyramidal cell cluster.

During data preprocessing, a central core region of 0.852 mm *×* 0.852 mm was selected for subsequent alignment (Fig. 1b). Based on the registered volume, we established a multi-resolution processing framework, to balance computational efficiency and structural fidelity. Nuclei were segmented at 128 nm resolution for large-scale cellular census (Fig. 1f). Neuronal somata, myelin, blood vessels, and coarse cellular morphologies were extracted at 32 nm resolution (Fig. 1f–g), whereas fine neuronal processes and synaptic contacts were reconstructed at 8 nm resolution.

Neuronal skeletons (Fig. 1h) are treated as the primary representation in neuronal proofreading. This choice is motivated by the presence of both split errors and merge errors in automated segmentation. Skeletons and meshes are therefore jointly utilized to reconstruct neurons. Unlike the segmentation-based proofreading workflow in CAVE, our sampled-skeleton strategy allowed early iterations of segmentation and skeleton-based proofreading to proceed in parallel. For some merge errors, segmentation rollback and patching techniques were used for correction.

Based on our dataset, we first focus on the analysis of pyramidal cell clusters, a key organizational structure in layer II of the MEC. Because L2P neurons are also distributed between clusters and no sharp boundary defines an L2P cluster, we used the internal exclusion core (ExC; described below) as the principal spatial reference for reconstruction. The ExC-centred circuit was completely reconstructed, including its constituent pyramidal cells and other neurons, together with a large population of neighboring neurons and portions of adjacent clusters, to delineate the local microcircuitry associated with the cluster. To our knowledge, this represents the first complete reconstruction and circuit-level analysis of a pyramidal cell cluster since the discovery of pyramidal cell aggregation in layer II of the MEC [15–17]. This reconstruction revealed strong coupling among cellular distribution, neuronal morphology, and synaptic connectivity, providing an integrated view of how cellular aggregation and regional segregation contribute to the organization of local circuit architecture.

Importantly, the value of this dataset extends beyond this focal circuit: it encompasses the MEC–PaS transition zone, the key upstream region of the MEC, together with all six MEC layers [18, 32]. The dataset therefore provides a rich resource for investigating the structural organization and connectivity of the MEC and its interactions with upstream circuits, with broad implications for understanding neural circuits underlying spatial navigation.

## PaS-MEC cell types and spatial organization

To characterize cell distributions in our dataset, we developed an automated image-based cell classification method (Fig. 2a). The method first detects nuclei at 128-nm resolution to localize individual cells and then classifies them using multi-view image patches of their cell bodies. Notably, the method requires only raw EM images as input, enabling the classification of most cells without requiring prior reconstruction of cellular morphology [33].

**Figure 2.**
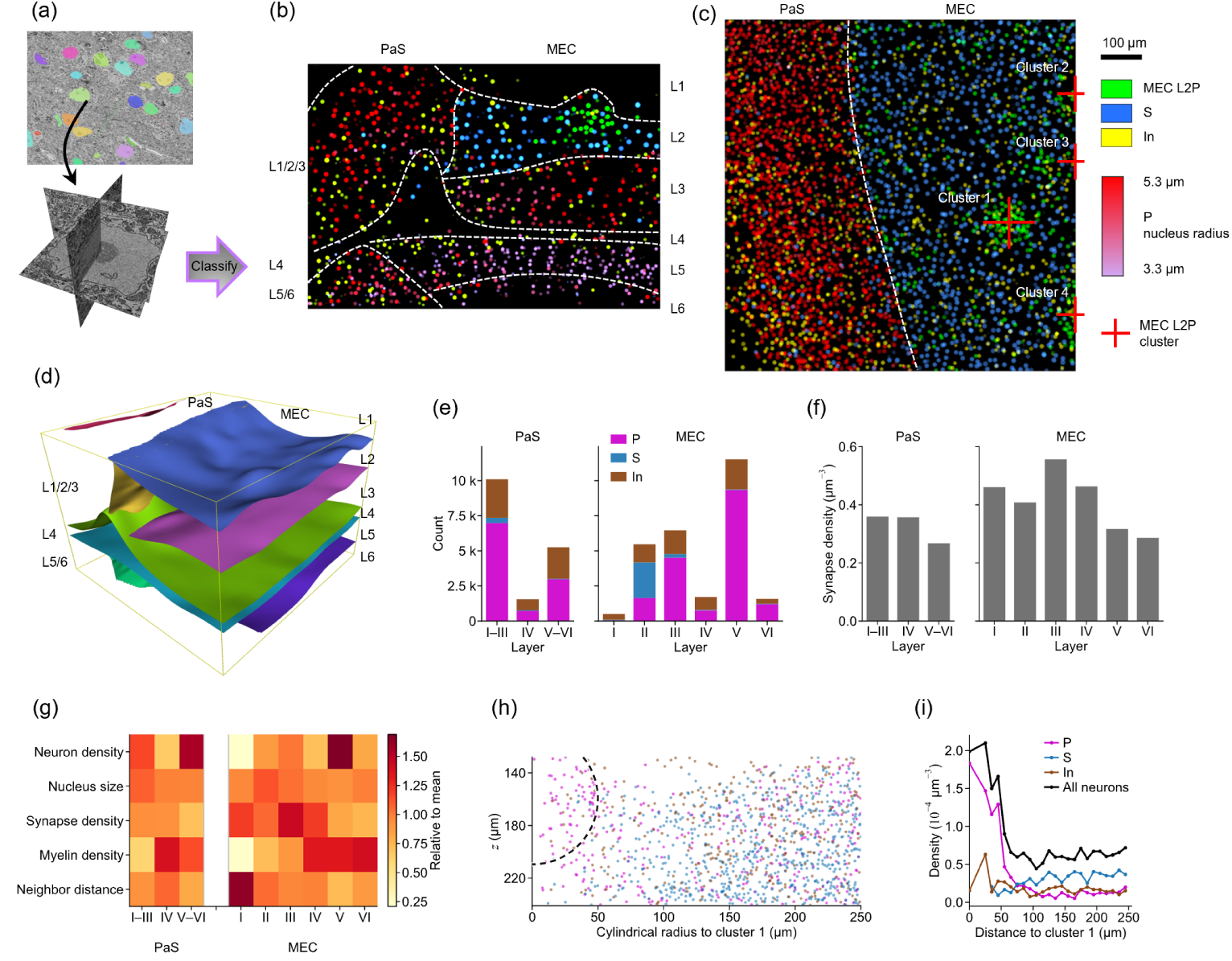
PaS - MEC layers and cell type distribution. (a) Schematic of the image-based artificial intelligence pipeline for automated localization and classification of cells. Nuclear segmentation was used to localize cells (top), and cell types were classified from multiple image sections passing through the soma centre (bottom). (b–c) Maps of pyramidal (P), stellate (S) and inhibitory (In) cell distributions in the horizontal plane (b), shown as a 10 *µ*m-thick projection along the *y* axis through L2P cluster 1, and in the tangential plane (c), shown as a 50 *µ*m-thick projection along the *z* axis through the cluster. Laminar boundaries in the MEC and PaS are indicated, revealing marked differences in cell-type distributions across these boundaries. (d) Three-dimensional rendering of the laminar boundaries in the MEC and PaS. (e) Stacked counts of classified neuronal nuclei across grouped PaS layers I–III, IV and V–VI and MEC layers I–VI. (f) Synapse density in the corresponding PaS and MEC laminar compartments. (g) Relative neuron density, mean nucleus size, synapse density, myelin density and mean nearest-neighbour soma distance across PaS and MEC layers; each metric is normalized to its mean across the anatomical compartments. (h) Cylindrical projection of layer II P, S and In somata around the centre of pyramidal cluster 1, with azimuth collapsed onto cylindrical radius and depth retained on the *z* axis. The arc marks a three-dimensional distance of 50 *µ*m from the cluster centre. (i) Radial density profiles of P, S, In and all neurons as a function of three-dimensional distance from the centre of cluster 1. Densities outside the central 25 *µ*m sphere were calculated from cell counts normalized by the sampled volume of each radial shell.

This classification identified multiple neuronal cell types, yielding a total of 38,334 neurons after excluding vascular and glial cells. We further characterized the spatial distribution of pyramidal cell subtypes based on the locations and nuclear features of individual pyramidal cells, together with the distribution of stellate cells. These cellular distributions enabled us to delineate the boundary between the MEC and PaS and to resolve the laminar organization of the MEC [18, 32] (Fig. 2b, c).

We quantified neuronal cell types and cellular features across the different layers and regions (Fig. 2e–g). MEC layer V had the highest neuronal density and the smallest mean nucleus size among the MEC layers, with chain-like arrangements of layer V cells visible in Fig. 2b. Synapse density was highest in MEC layer III and declined progressively from layers IV to VI (Fig. 2f). Because the internal boundaries within PaS layers I–III and V–VI were not clearly resolved, these layers were analysed as grouped compartments; PaS layers I–III combined a relatively high neuronal density with a large mean nucleus size (Fig. 2g). In both the PaS and MEC, myelin density was higher in layers IV–VI than in the superficial layers. Although MEC layer II had neither the highest neuronal density nor the highest synapse density, its neurons had relatively large nuclei and exhibited the most conspicuous spatial organization: four clusters of pyramidal cells were identified (Fig. 2c), of which cluster 1 was fully contained within the volume.

Pyramidal cluster 1 comprised predominantly L2Ps, accompanied by a smaller population of interneurons (In) and very few stellate cells (S) (Fig. 2h, i). L2P somata formed a compact patch within the layer II band, consistent with the canonical MEC layer II lattice described previously in rodents [15, 17]. Although L2Ps were also present beyond the cluster, radial density analysis around cluster 1 revealed a sharp decline in L2P density at a radius of approximately 50 *µ*m (Fig. 2h, i). Beyond this radius, stellate-cell density gradually increased as L2P density declined, with stellate cells intercalated around the patch rather than concentrated within its core, consistent with previous anatomical and physiological observations [2, 11, 34].

Proofread reconstructions covered the cluster and nearby neurons for local connectivity analysis. Together, these data reveal an excitatory pyramidal patch embedded within a surrounding stellate- and interneuron-rich matrix.

## L2P apical dendrites predominantly extend dorsal-laterally, markedly tilted to the laminar structure

Observation of reconstructed layer 2 pyramidal (L2P) neurons revealed that MEC layer 2 pyramidal neurons extend an apical dendrite that is markedly tilted relative to the MEC laminar plane (Fig. 3a–c). By defining the apical vector as the vector from the soma toward a point proximal to the apical bifurcation, quantitative analysis showed that this vector deviated substantially from the canonical pia-directed axis in MEC layer 2 pyramidal neurons (Fig. 3c–f). This deviation was larger in L2P Cluster 1 than in other L2P neurons (Fig. 3e, f) [15].

**Figure 3.**
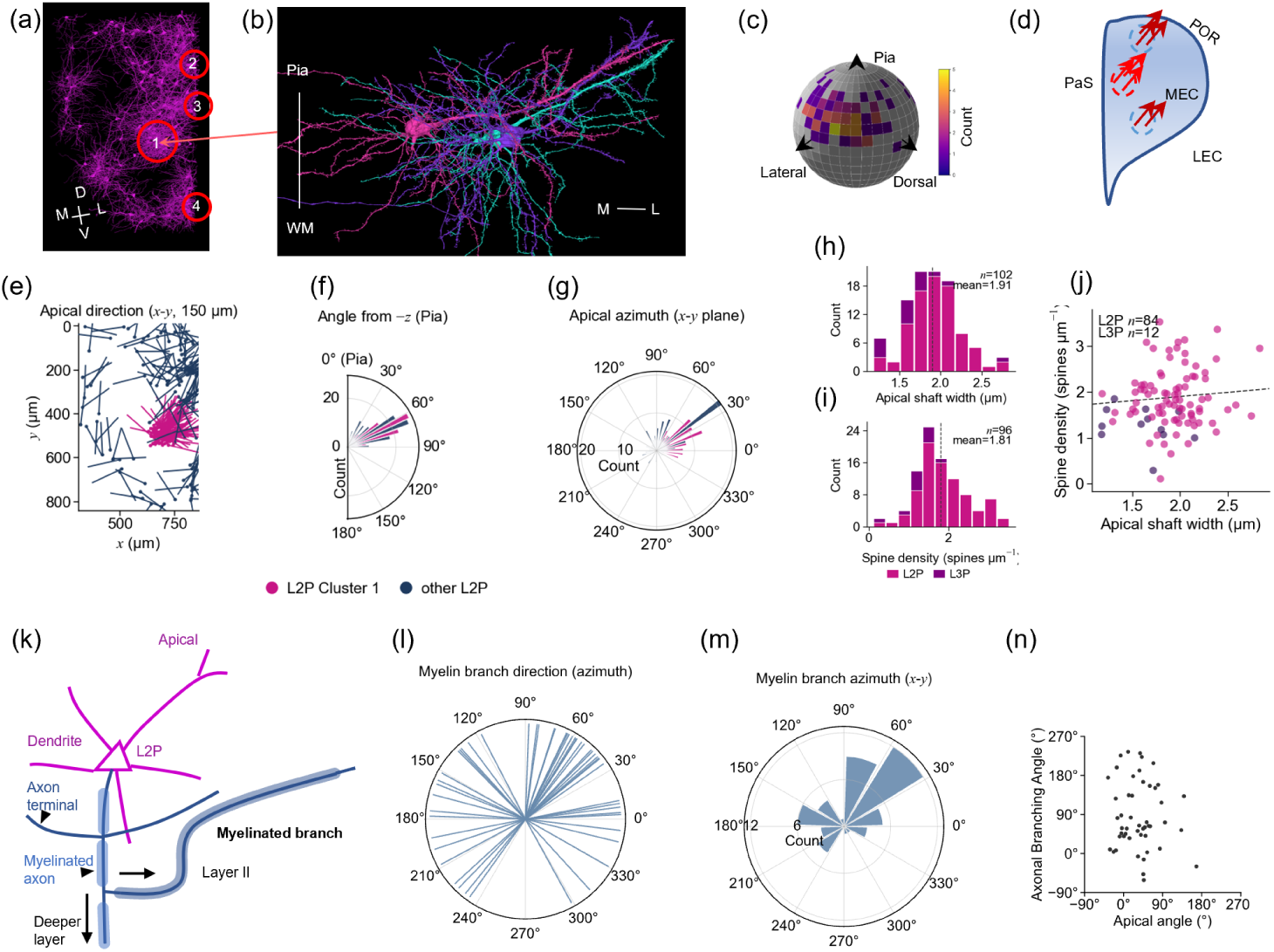
Morphology of apical dendrites and axonal branches of L2P neurons in MEC layer II. (a) Renderings of proofread L2P neurons in MEC layer II (120 representative neurons shown), with MEC L2P clusters identified in red circle. (b) Three pyramidal neurons within the first pyramidal cell cluster, showing apical dendrites with large deviations from the pia orientation. (c) Spherical distribution of apical dendrite orientations, revealing a predominant lateral-dorsal preference. (d) Schematic illustration of apical dendrite orientation. Because dendrites receive synaptic inputs, arrows indicate the direction from the apical dendritic tip toward the soma. (e–g) Apical dendrite orientation vectors and corresponding soma distributions (e), angular distribution relative to the pia-directed axis (f), and azimuthal angle distribution in the *xy* plane (g). The mean angle from the canonical pia-directed axis was 46.52 *±* 23.92*^◦^* (*n* = 214); L2P Cluster 1 showed a larger deviation (59.42 *±* 19.32*^◦^*, *n* = 88) than other L2P neurons (36.25 *±* 23.31*^◦^*, *n* = 126). In the *xy* plane, Cluster 1 apical dendrites had a mean azimuth of 31.28 *±* 36.98*^◦^* (*n* = 88). Across all reconstructed L2P neurons, most apical dendrites were oriented dorsolaterally (0*^◦^*–90*^◦^*, 63.1%, *n* = 135), with smaller fractions toward the dorsomedial (90*^◦^*–180*^◦^*, 14.5%, *n* = 31) and ventrolateral (270*^◦^*– 360*^◦^*, 14.5%, *n* = 31) axes. (h–j) Apical dendritic shaft width (h) and spine density (i) of L2P neurons in Cluster 1 and nearby L3P neurons. (j) Scatter plot of shaft width versus spine density; points form a continuous distribution without separable morphological subclasses within Cluster 1. (k–m) Myelinated axonal branches of L2P neurons extending within MEC layer II (k), normalized branch orientation vectors (l), and directional distribution analysis (m). Dorsolateral, ventrolateral, and medial directions accounted for 53.1%, 10.2%, and 36.7%, respectively. (n) Apical azimuth versus axonal branching azimuth in the *xy* plane for L2P neurons with both measurements (*n* = 49; 0*^◦^* = +*x*, 90*^◦^* = *−y*; same convention as in (g, m)). Each point is one neuron. The two orientations showed no evidence of statistical dependence (permutation test, *P* = 0.91).

Analysis of apical dendritic orientation in the *xy* plane further revealed a bias toward the dorsolateral direction in L2P Cluster 1 and across all reconstructed L2P neurons in the MEC (Fig. 3g), although this preference was not absolute, as a substantial fraction of dendrites were oriented toward the dorsomedial and ventrolateral axes. In the MEC, the dorsolateral, dorsomedial and ventrolateral directions are anatomically adjacent to the postrhinal cortex (POR), PaS and lateral entorhinal cortex (LEC), respectively [18, 32, 35]. Apical dendrites of MEC L2P neurons exhibited a dorsolateral bias, contrasting with the medial location of the upstream PaS [18].

Previous work has subdivided layer II pyramidal neurons into distinct subclasses [7, 12]. Although apical dendritic shaft width and spine density varied across L2P neurons in cluster 1 (Fig. 3h–i), clustering on these morphological features did not resolve separable subclasses within the L2P population (Fig. 3j).

Many L2P neurons also extended a myelinated axonal branch within MEC layer II, in some cases for more than 500 *µ*m. Statistical analysis of branch orientation revealed heterogeneity across cells: some myelinated branches preferentially extended dorsolaterally, whereas others projected medially (Fig. 3k–m). Although both myelinated-branch and apical-dendrite azimuths clustered toward the dorsolateral direction, the two orientations showed no significant association across cells (permutation test, P=0.91, n=49; Fig. 3n), suggesting no evidence of statistical dependence.

Together, apical dendrites and myelinated axonal branches represent the main long-range processes of L2P neurons within layer II, indicating that these cells engage in extensive intra-laminar wiring rather than relying solely on short-range local contacts mediated predominantly by basal dendrites.

## A central exclusion core shapes neural morphology

Following observations of L2P apical dendrites and myelinated axon-branch, we found that, in addition to the previously reported phenomenon of layer III pyramidal neuron (L3P) apical dendrites circumventing pyramidal cell clusters [15, 17], dendrites and axons from nearby excitatory stellate cells, pyramidal neurons, and some inhibitory interneurons also exhibit trajectories that bypass the pyramidal cluster (Fig. 4a, b) [16, 34]. By reconstructing the inner envelope of multiple external neuronal morphologies around L2P cluster 1, **an exclusion core** (ExC) at the center of L2P cluster is discovered, which is well approximated by an ellipsoid.

**Figure 4.**
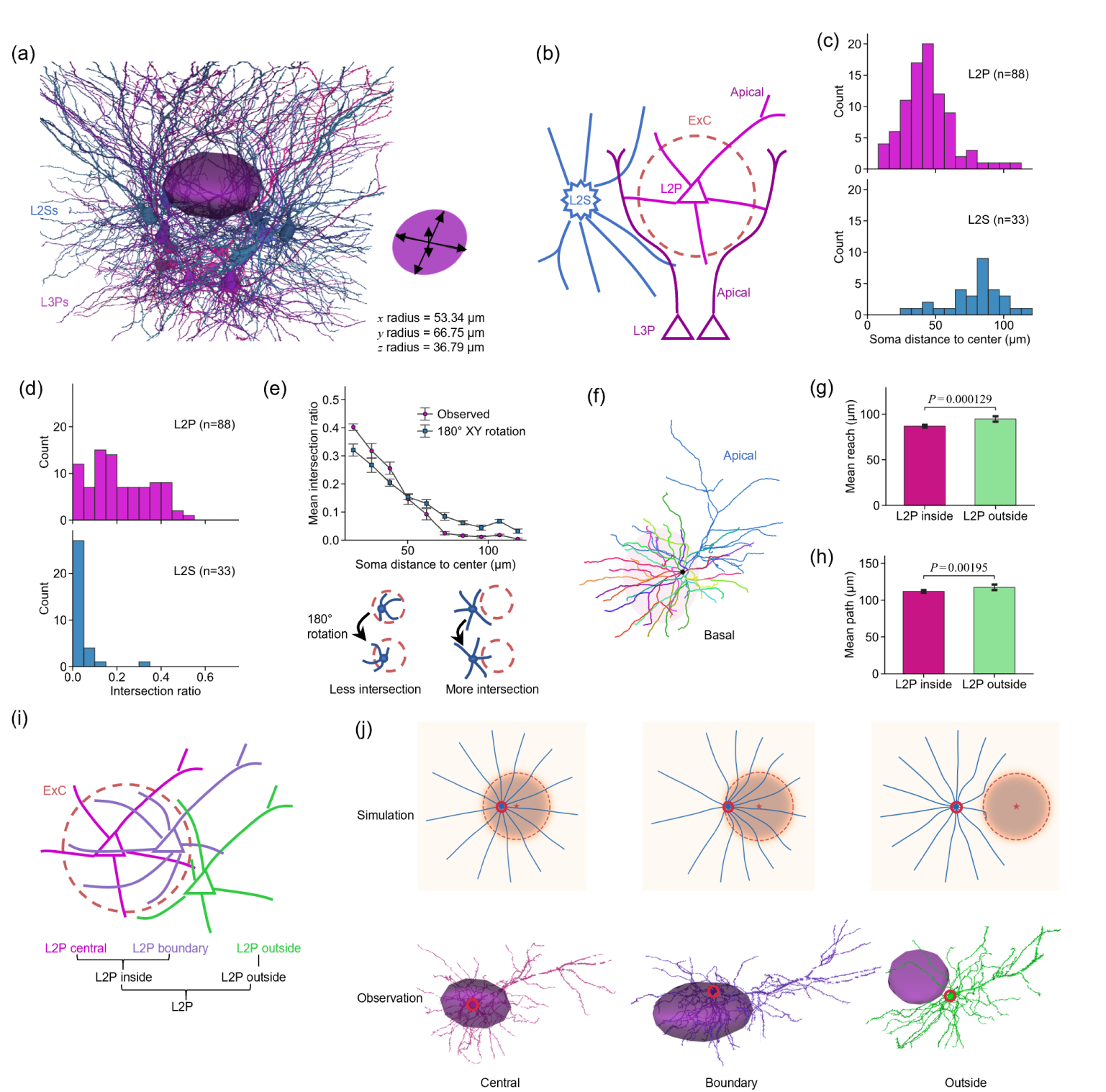
An exclusion core (ExC) at the center of pyramidal cell clusters. (a) Renderings of the exclusion core and surrounding L2S and L3P dendrites. The exclusion core located at the center of the pyramidal cell cluster was approximated by an ellipsoid. Nearby neurons, including L2S and L3P cells, exhibited dendritic trajectories that circumvented this core. (b) Schematic illustration of the influence of the exclusion core on neighboring neuronal morphology. (c) Distribution of soma distances from the exclusion core center for nearby L2P and L2S neurons. (d) Distribution of dendritic intersection ratios with the exclusion core for nearby L2P and L2S neurons. (e) Relationship between dendritic intersection ratio and soma distance from the exclusion core center for nearby L2P and L2S neurons. A control experiment rotating neurons by 180*^◦^* around the *z* axis in the *xy* plane revealed a distance-dependent bias: neurons within 50 *µ*m exhibited a tendency toward the exclusion core center, whereas neurons beyond 50 *µ*m preferentially grew away from the exclusion core. (f) Schematic illustration of dendritic compartmentalization in pyramidal neurons. For the basal dendrite quantification shown in (g,h), apical dendrites were excluded, while basal dendrites of all pyramidal neurons were separately quantified according to their terminal branches. (g) Mean basal terminal reach (Euclidean terminal–soma distance) for L2P neurons inside (*n* = 38) versus outside (*n* = 51) the exclusion core. Bars show mean *±* s.e.m. (inside 86.7 *±* 1.5 *µ*m; outside 94.7 *±* 3.0 *µ*m; Mann–Whitney *P* = 1.3 *×* 10*^−^*^4^). (h) Mean basal terminal path length (skeleton distance from terminal to soma) for the same inside versus outside L2P cohorts. Bars show mean *±* s.e.m. (inside 111.8 *±* 1.9 *µ*m; outside 117.4 *±* 3.7 *µ*m; Mann–Whitney *P* = 1.9 *×* 10*^−^*^3^). (i) Schematic illustration of basal dendrite morphologies influenced by the exclusion core. Central L2P neurons extended radially outward through the exclusion core, boundary L21P2 neurons exhibited asymmetric deformation, and outside L2P neurons displayed circumferential dendritic arrangements around the exclusion core. (j) Simulation and experimental comparison of exclusion-core effects on basal dendrite growth. Central L2P neurons extended through the exclusion core; boundary L2P neurons showed preferential growth toward the exclusion core and increased deformation relative to the opposite side; outside L2P neurons formed circumferential dendritic trajectories that terminated at the exclusion-core boundary.

We then quantified the overlap ratio between individual neurons and the exclusion core, defined as the fraction of total basal dendritic length located within the fitted ellipsoid (Fig. 4c, d). We found that the exclusion core is strongly enriched for pyramidal neurons, whereas stellate cells show a substantially lower overlap probability (Fig. 4d). Notably, even neurons located in close proximity to the exclusion core centroid (i.e., with soma-to-centroid distances much smaller than their mean dendritic length) often exhibit minimal intersection with the core, suggesting a robust spatial avoidance phenomenon (Fig. 4e).

To test whether dendritic structures are spatially organized relative to the exclusion core, we performed a rotation-based control analysis. Each neuron was rotated by 180*^◦^* around the z-axis passing through its soma while preserving its intrinsic dendritic geometry. Following rotation, neurons near the core centre showed a reduced overlap with the exclusion core, whereas neurons farther from the centre exhibited an increased overlap (Fig. 4e). These opposing trends changed rapidly at around 50 *µ*m from the core centre, providing an approximate spatial scale for distinguishing neurons with dendrites preferentially located within the core from those whose dendrites were biased away from it. Notably, without prespecifying the boundaries of the exclusion core, this perturbation-based analysis revealed a spatial association between neuronal location and dendritic organization, providing independent support for the presence of an exclusion core at the centre of the pyramidal cell cluster.

To test our hypothesis that the exclusion core exerts a suppressive effect on dendritic growth, we compared basal dendritic extent between L2P neurons located inside versus outside the exclusion core. For each neuron, after excluding the apical tree, we computed the mean Euclidean reach and mean path length of its basal terminals (Fig. 4g, h). Inside L2P neurons exhibited significantly shorter mean terminal reach and mean path length than outside L2P neurons. This suggests that the reduced basal extent is consistent with spatial constraints on dendritic growth within the ExC. Consistent with this interpretation, prior anatomical work shows that reelin-positive stellate cells and related neurites avoid the calbindin-rich pyramidal island [8, 16, 34]. Intriguingly, a subset of pyramidal neurons also distributed away from the exclusion core (Fig. 4c), indicating that not all pyramidal somata are contained within the ExC boundary. This implies that the exclusion core is spatially nested within a broader pyramidal cell cluster, whose somatic extent exceeds the dendritically defined exclusion boundary.

To further characterize the morphology of L2P neurons relative to the exclusion core, we compared pyramidal neurons located inside, at the boundary of, and outside the exclusion core (Fig. 4i).

### Inside the exclusion region

Neurons located within the exclusion core exhibited relatively symmetric dendritic morphologies and extended processes that crossed the exclusion boundary.

### At the boundary of the exclusion region

Neurons positioned at the boundary but still within the exclusion core displayed a pronounced tendency for dendrites to grow preferentially along the internal side of the boundary, often forming arc-like trajectories that follow the exclusion interface before eventually crossing it at more distal segments.

### Outside the exclusion region

In contrast, neurons located outside the exclusion core tended to avoid the boundary altogether, with dendritic branches either circumventing the exclusion core or appearing to be reflected away from it.

Based on the characteristics of the exclusion core, we implemented a computational model of dendritic *growth* to investigate how dendritic morphogenesis arises under the influence of the exclusion field (Fig. 4j). In this model, each soma emits dendritic branches that extend in discrete steps. For simplicity, apical dendrites were omitted to focus on basal dendritic growth and its interaction with spatial constraints. Dendrite extension is governed by a gradient-driven process combining a sigmoidal exclusion potential field with mutual repulsion between neighboring dendrites. This model qualitatively recapitulates the observed phenotypes, including dendritic penetration within the exclusion core, boundary-following trajectories, and avoidance of the exclusion core by external neurons. Notably, the model recapitulates the counterintuitive dendritic distribution observed in boundary neurons: even when dendritic growth is initialized uniformly across all directions, the resulting morphology contains a higher density of dendrites within the exclusion core due to the influence of the exclusion field. Beyond reproducing the observed characteristics of dendritic growth, a key implication of the model is that the exclusion potential is best described by a sigmoid-like potential field, which varies gradually within the interior of the exclusion core and decays rapidly toward zero near its boundary.

## The central exclusion core shapes local connectivity

To examine how the pyramidal cell cluster and its exclusion core shape connectivity, we reconstructed dendrites and axons of 97 neurons in L2P cluster 1 and assembled a synaptic connectivity matrix for this layer II microcircuit with L2P inside and outside the exclusion core of cluster 1 distinguished (Fig. 5a). Connectivity analysis revealed that the exclusion core strongly structured nearby wiring. The most prominent effect was a compartment-specific difference in pyramidal connectivity across the exclusion boundary. Among interneurons, basket cells (BCs) and Re-MC neurons formed distinct cell-type-specific connection motifs (Fig. 5b).

**Figure 5.**
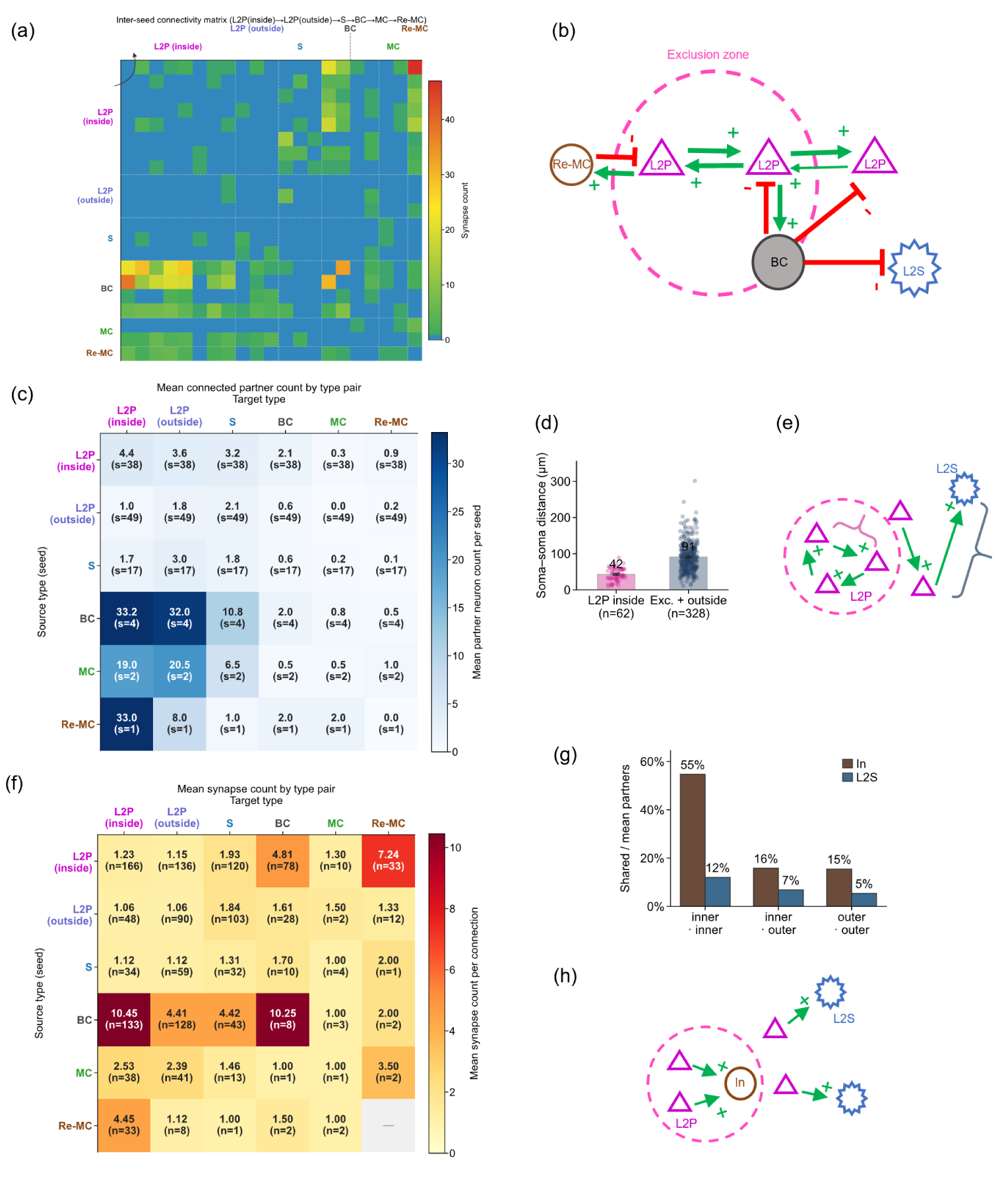
Cell-type-specific connectivity within pyramidal cell cluster. (a) Connectivity matrix showing synaptic connections among neurons within pyramidal cell clusters. BC denotes basket cells; Re-MC denotes recurrent Martinotti cells participating in recurrent inhibitory loops with layer II pyramidal neurons (L2P) within the same cluster. (b) Schematic representation of the cell-type-specific connectivity architecture related to pyramidal cell cluster. (c) Average number of postsynaptic partners between neuronal types. The number of seeds (*s*) represents the number of axon-containing neurons used for calculating partner statistics. (d) Soma-to-soma distances between seed and target neurons. Mean distances were *<* 42 *µ*m between L2Ps inside the exclusion core, less than half the 91 *µ*m distance between excitatory neurons outside the core. (e) Schematic illustration showing that connections within the exclusion core have shorter seed-to-target soma distances than those outside the exclusion core, indicating tighter local connectivity. (f) Average number of synapses per connection between neuronal types. *n* represents the number of neuron pairs with identified synaptic connections included in the analysis. (g) Shared downstream inhibitory and stellate-cell partners of layer II pyramidal neurons (L2P) loca_1_t_5_ed inside (inner) and outside (outer) the exclusion core of pyramidal cell clusters. (h) Schematic illustration of shared downstream connectivity patterns of L2P neurons inside and outside the exclusion core.

### Pyramidal neurons are more densely interconnected within the exclusion core than outside

Inside L2P neurons formed more connections with one another than they received from outside the exclusion core. On average, each inside L2P neuron connected to 0.83 other inside L2P neurons, whereas outside L2P neurons and stellate (S) cells connected to only 0.36 and 0.30 inside L2P partners, respectively (Fig. 5c). Because all inside L2P neurons in cluster 1 were fully reconstructed, incoming connectivity onto inside L2P cells is complete in our sample. Connectivity onto outside L2P and S neurons is likely underestimated, as these populations were only partially reconstructed. Even with this conservative sampling, excitatory connections originating outside the exclusion core exceeded the number of connections onto inside L2P neurons. Nevertheless, inside L2P pairs were linked at substantially shorter soma–soma distances (mean 42.4 *µ*m) than excitatory connections involving outside neurons (mean 90.7 *µ*m), indicating that the exclusion core supports spatially compact excitatory coupling (Fig. 5d–e) [11, 13].

We used the number of synaptic contacts as a proxy for connection strength. For each connected pre- and postsynaptic pair, we report the mean number of synaptic contacts per connected pair (hereafter, *synapses per pair*; Fig. 5f). Excitatory connections, including those among inside L2P neurons, typically carried only a few synapses per pair (L2P–L2P: 1.0–1.60; L2P to S: 1.36 and 1.95 for inside and outside compartments, respectively; Fig. 5f) [11, 12]. Thus, the cluster forms a dense, spatially confined excitatory subnetwork with individually weak synaptic connections. When considered as a whole, the L2P population within the ExC may avoid excessive pairwise coupling, as stronger coupling could promote more tightly coordinated and similar firing patterns and constrain population-level diversity. The observed architecture of ExC may instead favour a rich repertoire of population activity within the patch.

### Interneurons and inside L2P neurons form a compartment-specific recurrent inhibitory microcircuit

In contrast to weak excitatory coupling, inhibitory neurons formed stronger contacts with L2P cells. BCs innervated inside L2P neurons with a mean of 12.50 synapses per pair, substantially exceeding BC-to-outside-L2P pairs (6.17 synapses per pair; Fig. 5f). Re-MC neurons contacted many inside L2P partners; their synaptic strength was lower than that of BC–L2P pairs but higher than excitatory L2P–L2P coupling (3.86 versus 1.23 synapses per pair).

Both BCs and Re-MC neurons participated in a recurrent inhibitory motif linking inside L2P neurons (L2P_inside_–In–L2P_inside_; Fig. 5a–d) [13, 14, 36, 37]. For both L2P_inside_*→*In and In*→*L2P_inside_ directions, mean synapses per pair exceeded values for other cell-type combinations (Fig. 5f), a pattern not observed for outside L2P neurons. This recurrent inhibitory microcircuit appears specific to pyramidal neurons and interneurons within the exclusion core.

The two interneuron classes implemented distinct versions of this motif. BC axons extended beyond the exclusion core and inhibited diverse excitatory and inhibitory targets, consistent with a patch-scale gating role. Re-MC neurons were more selective, targeting mainly inside L2P apical dendrites. We propose that BCs mediate intercompartment suppression, whereas Re-MC cells provide intra-core feedback control within the broader MEC layer II circuit [38–40].

### Pyramidal neurons share interneuron partners more than stellate partners

Because L2P axons arborize widely across layer II, we asked whether neighbouring pyramidal cells converge onto shared postsynaptic targets. Pairwise analysis of downstream partner overlap showed that L2P neurons inside the exclusion core shared interneuron partners more often than stellate partners (Fig. 5g–h) [8, 40]. The L2P share of L2S partners was low both inside and outside the exclusion core. This pattern is consistent with a bias toward co-innervation of local inhibitory interneurons by nearby pyramidal cells.

Together, these analyses indicate that the exclusion core organizes layer II pyramidal cluster connectivity—promoting dense excitatory coupling within the core and embedding a recurrent inhibitory microcircuit in which BCs may serve as a key hub for compartment-level control.

## Inhibitory recurrent connectivity in the pyramidal cell cluster

Inhibitory neurons can exert distinct functional effects depending on the subcellular location of their postsynaptic targets [38, 39]. For example, Martinotti cells (MCs) preferentially target apical dendrites, basket cells (BCs) target somata, and chandelier cells target axon initial segments, and may therefore regulate synaptic input, spike generation, or neuronal output, respectively [36, 37, 41, 42]. Within the recurrent inhibitory circuit of the pyramidal cell cluster, we identified two distinct forms of inhibitory feedback: BCs primarily target the perisomatic region, whereas Re-MC neurons preferentially target the apical dendritic shaft.

### Recurrent Martinotti cells targeting apical dendrites of L2Ps in the exclusion core

The dendrites of Re-MC neurons extend into the pyramidal cell cluster, whereas their axons course around the upper boundary of the cluster into layer 1, where they contact the apical dendrites of L2P neurons (Fig. 6a–c). Because the apical dendrites of L2Ps are markedly tilted and predominantly oriented toward the dorsolateral (DL) direction, the axons of Re-MC neurons similarly extend toward the DL direction and form contacts with L2P apical dendrites.

**Figure 6.**
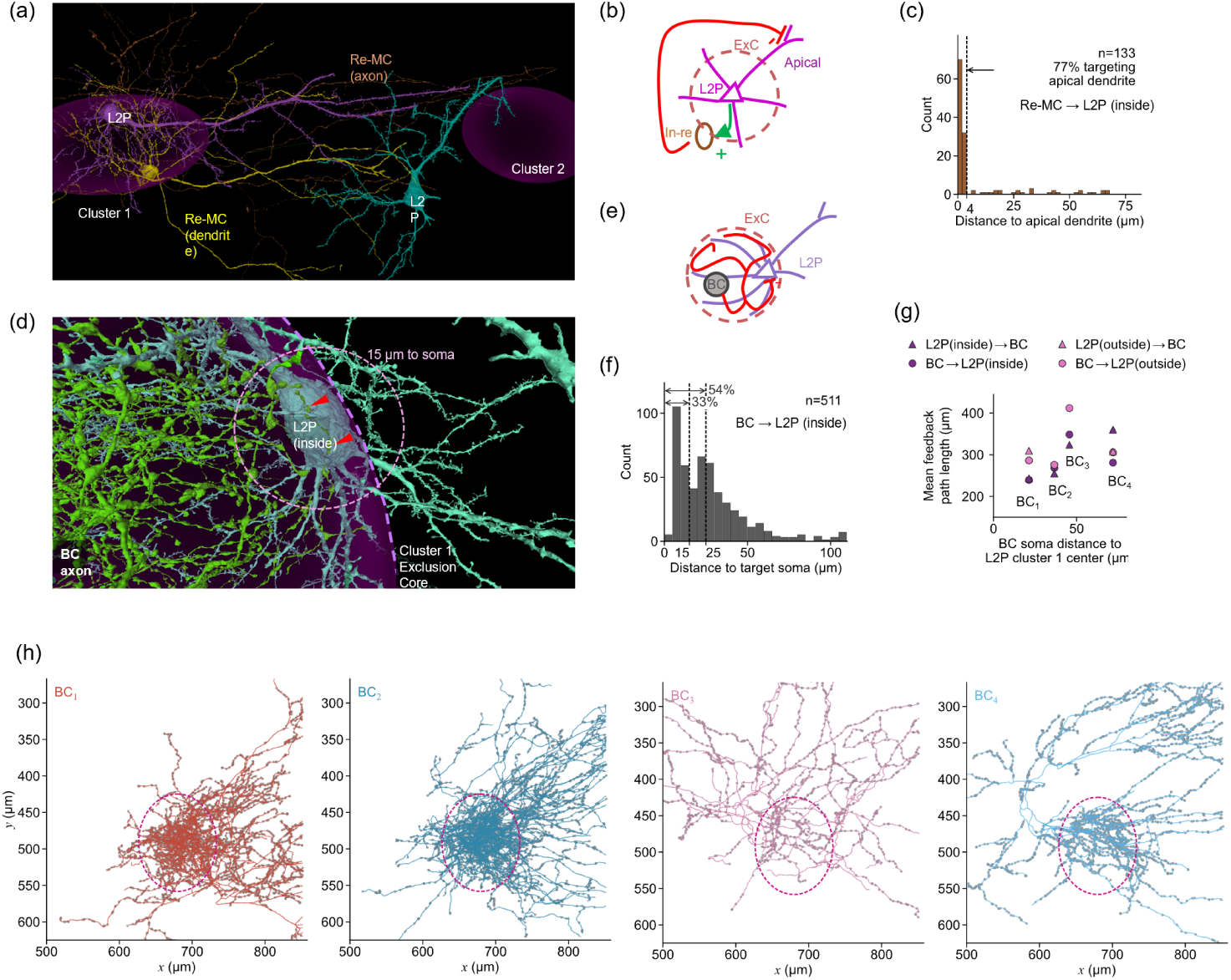
Distinct inhibitory feedback circuits mediated by recurrent Martinotti cells (Re-MC) and basket cells within pyramidal cell clusters. Re-MC neurons target the apical dendrites of layer II pyramidal neurons (L2P), whereas basket cells (BCs) target the somatic and perisomatic regions of L2P neurons within the exclusion core. (a) Example of Re-MC neurons targeting L2P apical dendrites. The reconstructed Re-MC neuron forms synaptic contacts with pyramidal neurons located both within the exclusion core of cluster 1 and outside the cluster. (b) Schematic illustration of the Re-MC-mediated inhibitory feedback circuit. (c) Distribution of distances from Re-MC synapses to the apical dendrites of target L2P neurons, showing that about 77% of Re-MC synapses were located on apical dendrites. (d) Basket cells form multiple synaptic contacts (red arrows) with L2P somata located within the exclusion core. The exclusion core (purple) is clearly visible, and basket-cell axonal terminals within this region exhibit highly tortuous trajectories that remain confined inside the exclusion core. (e) Distribution of distances from BC synapses to the soma centres of target L2P neurons within the exclusion core, showing that approximately 33% of BC synapses were located in the perisomatic region. (f) Schematic illustration of basket-cell targeting of the somatic and perisomatic regions of L2P neurons within the exclusion core. (g) Mean L2P–BC feedback path length versus BC soma distance to cluster 1 centre (inside vs. outside L2P partners; triangles, L2P*→*BC; circles, BC*→*L2P). (h) *xy* projections of BC axons and synapses (BC_1_–BC_4_ colour-coded by proximity to cluster centre; purple dashed ellipse, ExC).

The involvement of Re-MC neurons in this recurrent inhibitory circuit suggests that they may regulate excitatory drive to the apical dendrites of L2Ps within the exclusion core. Specifically, their preferential targeting of apical dendritic shafts may provide a mechanism for suppressing dendritic inputs when neurons within the pyramidal cell cluster become strongly active, consistent with Martinotti-cell control of apical input [36].

### Basket cells forming recurrent inhibitory circuits targeting perisomatic regions of L2Ps in the exclusion core

A subset of BC axons was densely distributed within the exclusion core (ExC) of the pyramidal-cell cluster (Fig. 6d). Rather than extending readily beyond the ExC boundary, these axonal arbors tended to remain within the core, forming dense, locally confined terminal fields. Their terminals broadly targeted the perisomatic regions of L2P neurons within the ExC (Fig. 6e, f) [8, 40]. This dense perisomatic innervation, together with the compact reciprocal connectivity (Fig. 6g) between L2P neurons and BCs, suggests that BC recruitment could provide rapid and spatially coordinated inhibition across the ExC, potentially narrowing the temporal window for pyramidal-cell firing and favouring synchronously structured activity.

Beyond the ExC, BC axons extended in oriented bands and formed synapses along their trajectories (Fig. 6h). Notably, the axons of all four reconstructed BCs extended towards cluster 3 along an azimuth of approximately 45*^◦^*. Synaptic terminals outside the ExC were concentrated in oriented bands, with azimuths measured from the ExC centre in the *xy* plane (0*^◦^* = +*x*, 90*^◦^* = *−y*) peaking at 37*^◦^* for BC_1_ and BC_2_, 92*^◦^* for BC_3_, and 58*^◦^* for BC_4_. These orientations were broadly aligned with directions towards neighbouring pyramidal-cluster territories, notably at approximately 37*^◦^*and 58*^◦^*. The projected full width of each band, estimated from the 10th–90th percentile envelope perpendicular to its peak azimuth, ranged from 35 to 52,*µ*m (mean, 42,*µ*m), consistent with a spatially confined strip approximately 50,*µ*m wide. Together, these anatomical features suggest that BCs may link local perisomatic inhibition within the ExC to spatially directed inhibitory output beyond the cluster, potentially extending the influence of cluster activity across layer II (Fig. 6h).

Together, Re-MCs and BCs provide distinct forms of recurrent inhibition, targeting apical dendrites and perisomatic regions of L2P neurons, respectively. Their complementary targeting patterns and the spatially extended BC axons suggest that the cluster may combine local regulation of dendritic input and pyramidal-cell firing with directionally organized inhibition across layer II.

## Discussion

We generated a synapse-resolution volume electron microscopy dataset spanning all six MEC layers, the adjacent PaS and their transition, enabling cellular organization and connectivity to be examined within a shared anatomical context. Within this dataset, our study reveals a spatially organized microcircuit centred on the layer II pyramidal-cell cluster. By combining a cellular census across the MEC–PaS volume with exhaustive reconstruction of one fully contained cluster, we identify the exclusion core as an organizing feature that links cell position, neurite geometry and synaptic connectivity. The ExC is not simply a region with fewer external neurites: it divides L2P neurons into spatial compartments with distinct basal dendritic architectures and wiring rules. Rotation controls and growth simulations further indicate that these patterns cannot be explained by soma position alone, but are consistent with a spatial constraint that acts during dendritic arborization. The recurrence of this organization in additional clusters suggests that the ExC is a general component of the pyramidal-patch architecture rather than an idiosyncrasy of cluster 1.

The reconstructed morphologies also reveal an unexpectedly strong long-range component within layer II. L2P apical dendrites were markedly tilted from the canonical pia-directed axis and preferentially oriented dorsolaterally, whereas many L2P neurons extended myelinated axonal branches for hundreds of micrometres within the layer. At the same time, their basal dendrites were organized locally around the ExC, with shorter arbors inside the core and circumferential or boundary-following trajectories outside it. These observations suggest that individual L2P neurons combine spatially constrained local integration through basal dendrites with longer-range sampling and output through apical dendrites and myelinated axonal branches. The absence of a cell-by-cell association between apical and myelinated-axon orientations further implies that these two long-range compartments need not participate in the same directional pathway.

This morphological partition was reflected in the synaptic network. Inside L2P neurons formed a dense and spatially compact, but individually weak, recurrent excitatory subnetwork and converged onto shared inhibitory partners. Feedback inhibition was subcellularly differentiated: BCs provided strong perisomatic inhibition, whereas the reconstructed Re-MC pathway preferentially targeted apical dendrites. Dense L2P–BC feedback provides a potential circuit mechanism for temporally coordinating pyramidal activity. The same BCs also extended beyond the ExC in oriented axonal bands, suggesting a route through which cluster-local activity could generate spatially directed inhibitory signals across layer II. Thus, the cluster may function not only as a compact local processing unit but also as a source of temporally structured inhibition at a broader regional scale.

The anatomical mosaic described here may nevertheless be relevant to the modular organization of grid-cell circuits. Pyramidal patches are arranged periodically and lie along the borders of grid modules [15, 19, 43], whereas continuous-attractor models commonly rely on relatively local excitation coupled to inhibition operating over a broader spatial range [4, 5]. The compact recurrent excitation within the ExC, shared recruitment of interneurons and outwardly directed BC projections provide an anatomical motif with these qualitative properties. In this view, each pyramidal cluster could support dense local processing while inhibitory outputs coordinate or separate activity across neighbouring territories. However, the present EM data do not identify the spatial firing phenotype of reconstructed neurons and therefore do not establish a one-to-one correspondence between an anatomical pyramidal cluster and a functional grid module. Instead, the connectivity identified here provides anatomical constraints that can be incorporated into future attractor models and tested against physiological measurements.

A complete mechanistic account of pyramidal-cluster function nevertheless remains out of reach. The contributions of other inhibitory cell types and the corresponding upstream and downstream signals cannot yet be resolved simultaneously. We therefore consider the more general architecture revealed here—spatially segregated domains supporting distributed computation while coupling to region-scale modulation—to be the more robust and transferable principle.

The dorsolateral bias of apical dendrites and the long myelinated axonal branches in layer II point to directional information flow beyond the immediate cluster, but morphology alone cannot determine the origins of dendritic input or the destinations and functional effects of these axons. Moreover, long processes are necessarily truncated at the boundaries of the EM volume. Resolving these pathways will require complementary modalities, including pathway-specific tracing, molecular identification of reconstructed cells, targeted electrophysiology and perturbation of candidate afferents from PaS, postrhinal cortex and other neighbouring regions. Such experiments will be important for determining whether the observed directional biases reflect shared regional pathways, cell-state-dependent routing or independent input and output organization.

The scope of the dataset extends beyond the cluster analysed here. Because the continuous volume includes all six MEC layers, the adjacent PaS and their transition, it offers an opportunity to reconstruct cross-layer and cross-regional pathways in a common anatomical coordinate system. Priorities include resolving the larger-scale stellate-cell network and its subtypes across layer II, tracing interactions between superficial and deep MEC layers, mapping PaS–MEC input–output relationships, and characterizing the myelinated zone at the MEC–PaS interface. The present connectivity analysis remains incomplete for neurons outside the fully reconstructed cluster, and conclusions based on rare interneuron types will require replication across additional cells and clusters. Extending proofreading and synapse identification across these domains should reveal which principles are cluster-specific and which generalize across the MEC–PaS circuit, providing a foundation for models that connect synaptic architecture to navigation-related population dynamics.

## Notes

### Competing Interest Statement

The authors have declared no competing interest.

### Summary of Updates

Only the author affiliations in the PDF were updated; no other changes were made.

